# Discovery of Novel Replication Proteins for Large Plasmids in Cyanobacteria and Their Potential Applications in Genetic Engineering

**DOI:** 10.1101/2023.10.09.561360

**Authors:** Kazuma Ohdate, Minori Sakata, Kaisei Maeda, Yutaka Sakamaki, Kaori Nimura-Matsune, Ryudo Ohbayashi, Wolfgang R. Hess, Satoru Watanabe

## Abstract

Numerous cyanobacteria capable of oxygenic photosynthesis possess multiple large plasmids exceeding 100 kbp in size. These plasmids are believed to have distinct replication and distribution mechanisms, as they coexist within cells without causing incompatibilities between plasmids. However, information on Rep proteins necessary for plasmid replication initiation in cyanobacteria is limited. *Synechocystis* sp. PCC 6803 hosts four large plasmids, pSYSM, pSYSX, pSYSA, and pSYSG, but Rep proteins for these plasmids, except for CyRepA1 on pSYSA, are unknown. Using Autonomous Replication sequencing (AR-seq), we identified two potential Rep genes in *Synechocystis* 6803, *slr6031* and *slr6090*, both located on pSYSX. The corresponding Rep candidates, Slr6031 and Slr6090, share structural similarities with Rep-associated proteins of other bacteria and homologs were also identified in various cyanobacteria. We observed autonomous replication activity for Slr6031 and Slr6090 in *Synechococcus elongatus* PCC 7942 by fusing their genes with a construct expressing GFP and introducing them via transformation. The *slr6031/slr6090*-containing plasmids exhibited lower copy numbers and instability in *Synechococcus* 7942 cells compared to the expression vector pYS. While recombination occurred in the case of *slr6090*, the engineered plasmid with *slr6031* coexisted with plasmids encoding CyRepA1 or Slr6090 in *Synechococcus* 7942 cells, indicating the compatibility of Slr6031 and Slr6090 with CyRepA1. Based on these results, we designated Slr6031 and Slr6090 as CyRepX1 (<u>Cy</u>anobacterial <u>Rep-</u>related protein encoded on pSYS<u>X</u>) and CyRepX2, respectively, demonstrating that pSYSX is a plasmid with “two Reps in one plasmid”. Furthermore, we determined the copy number and stability of plasmids with cyanobacterial Reps in *Synechococcus* 7942 and *Synechocystis* 6803 to elucidate their potential applications. The novel properties of CyRepX1 and 2, as revealed by this study, hold promise for the development of innovative genetic engineering tools in cyanobacteria.

## 1 Introduction

Cyanobacteria, a monophyletic lineage of Gram-negative oxygenic photosynthetic bacteria, thrive on Earth for at least 2.8 billion years, and have shaped the geochemistry and the biological landscape of the Earth (Garcia-Pichel et al. 2020). They occur ubiquitously as long as there is light, and inhabit a wide range of ecological conditions, from aquatic environments such as lakes, rivers, and oceans, to arid deserts, polar regions, caves, and even in symbiosis with other organisms such as fungi in the formation of lichens (Shih et al. 2013). Cyanobacteria have a physiology based on oxygen-producing photosynthesis, allowing them to produce biomass by using solar energy. Consequently, they have been recognized as the evolutionary ancestors of chloroplasts, the energy-producing organelles of plants and algae. Finally, cyanobacteria have recently gained attention for their potential as green cell factories for the CO2-neutral biosynthesis of various products (Jodlbauer et al. 2021, Knoot et al. 2018).

As a diverse group of organisms, cyanobacteria vary greatly in the size and structure of their genomes, which are composed of chromosomes and plasmids. While certain marine cyanobacteria, like *Prochlorococcus* and *Synechococcus*, exhibit compact genomes in size and lack plasmids, many other cyanobacteria, mainly inhabiting freshwater and terrestrial environments, show more intricate genome organization. Complete genome information is available in the KEGG database for 145 cyanobacteria (https://www.genome.jp/), 69 of which have one or more extrachromosomal elements and 43 of these harbors large plasmids over 100 kbp (Supplementary Data 1). For example, in *Synechocystis* sp. PCC 6803, the most widely studied model cyanobacterium, there are four large plasmids (pSYSM: 120 kbp, pSYSX: 106 kbp, pSYSA: 103 kbp, and pSYSG: 44 kbp) and three smaller plasmids (pCA2.4: 2.4 kbp, pCB2.4: 2.4 kbp and pCC5.2: 5.2 kbp) (Kaneko et al. 2003, Trautmann et al. 2012, Xu and McFadden 1997, Yang and McFadden 1993, Yang and McFadden 1994). In the context of plasmids, identical replication mechanisms lead to competitive interactions, resulting in incompatibility. This phenomenon, known as plasmid incompatibility, refers to the inability of plasmids with similar replication and partitioning mechanisms to coexist in the same host cell line. It has long served as an indicator for classifying plasmid types (Shintani et al. 2015). However, in certain cyanobacteria such as *Synechocystis* 6803, multiple plasmids are maintained simultaneously without incompatibility, suggesting that these cyanobacteria possess different replication mechanisms for maintaining multiple plasmids.

In *Synechocystis* 6803, Rep proteins, the factors involved in the replication initiation, have been identified in the smaller-sized plasmids, pCA2.4, pCB2.4, and pCC5.2. These plasmids are abundant per cell and are predicted to employ a rolling circle replication mechanism (Berla and Pakrasi 2012, Xu and McFadden 1997, Yang and McFadden 1993, Yang and McFadden 1994). Due to their high copy numbers, these plasmids have been widely employed as scaffolds of expression vectors, facilitating the production and development of valuable compounds (Jin et al. 2018, Opel et al. 2022, Sakamaki et al. 2023a, Sakamaki et al. 2023b). Notably, the Rep protein CyRepA2, encoded in pCC5.2, is highly conserved in various cyanobacteria. Consequently, we have successfully developed an expression vector pYS that is compatible with in a wide range of species (Sakamaki et al. 2023a). In contrast, our understanding of the replication mechanisms of the larger plasmids of *Synechocystis* 6803, specifically pSYSM, pSYSX, pSYSA, and pSYSG, remains limited, except for insights into pSYSA. It’s worth noting that a recent study has revealed an autonomous replication region within pSYSA that contains both the Rep protein and its regulatory domain (Kaltenbrunner et al. 2023). Rep protein homologs, designated CyRepA1 in case of pSYSA, are found in a wide spectrum of cyanobacteria, similar to observations for CyRepA2 (Sakamaki et al. 2023a). The regulatory region (*ssr7036*) immediately upstream of the *CyRepA1* gene (*slr7037*) is transcribed into sense and antisense small RNAs and likely plays a role in initiating θ-type replication (Kaltenbrunner et al. 2023). Notably, artificial plasmids VIII23, harboring *CyRepA1* and its regulatory region, was maintained not only in the original host, *Synechocystis* 6803, but also in *Synechococcus* 7942, indicating replication activity of CyRepA1 in several cyanobacterial hosts, akin to CyRepA2.

No genes bearing homology to CyRepA1 or other established replication proteins in cyanobacteria have been identified thus far for the other large plasmids in *Synechocystis* 6803, nor is there any insight into autonomous replication regions. This suggests the existence of unidentified mechanisms for replication initiation. In this study, aiming to identify novel Rep proteins in *Synechocystis* 6803, we conducted a library screening and found two candidate proteins for Rep, Slr6031 and Slr6090, both encoded in plasmid pSYSX. Homologs were found in various cyanobacteria with low sequence identity, while structural similarity to corresponding protein domains supported their replication-related functions. We constructed plasmids with candidate genes *slr6031* and *slr6090*, and tested their compatibility with other plasmids encoding other known Reps. Our results advance the understanding of enigmatic Rep proteins in cyanobacteria and facilitated their utilization in synthetic biology approaches involving cyanobacteria.

## 2 Material and Methods

### 2.1 Cyanobacteria strains and growth condition

This study used freshwater cyanobacteria *Synechococcus elongatus* PCC 7942 (our laboratory strain *S*. 7942 TUA) (Watanabe et al. 2012) and *Synechocystis* sp. PCC 6803 (substrain *S*. 6803 GT-I) (Kanesaki et al. 2012). Both strains were cultured in modified BG-11 medium containing twice the usual amount of sodium nitrate (final concentration 35.3 mM) and 20 mM HEPES-KOH (pH 8.2). Cultures were grown photoautotrophically at 30 °C, using continuous illumination (50 μmol photons m^-2^s^-1^) and with 2% CO2 (v/v) bubbling. When appropriate, chloramphenicol (Cm, 10 μg/mL), kanamycin (Km, 10 μg/mL), and spectinomycin (Sp, 40 μg/mL) were added to media.

### 2.2 Autonomous replication sequencing (AR-seq)

AR-seq was performed according to a previous study conducted to determine the region responsible for *Synechocystis* 6803 replication (Figure 1A) (Sakamaki et al. 2023a). A genomic library constructed by ligating 1.5-2.5 kbp fragments of *Synechocystis* 6803 genomic DNA with the ColE1 origin and *cat* gene (Library A) was transformed into *Synechocystis* 6803 cells to obtain 6.3 × 10^3^ colonies (Library B). To obtain plasmids that replicated independently of chromosomal integration, the DNA extracted from library B was introduced into *E. coli* again, and 456 transformants were obtained (Library C). Genomic libraries (Libraries A–C) were subjected to comprehensive sequencing analysis as previously described (Sakamaki et al. 2023a). To identify the regions included in the library, trimmed reads were mapped to the *Synechocystis* 6803 genome (accession numbers, chromosome: AP012276, pSYSM: AP004310, pSYSX: AP006585, pSYSA: AP004311, pSYSG: AP004312, pCA2.4: CP003270, pCB2.4: CP003271, and pCC5.2: CP003272) using the CLC Genomics Workbench ver. 20.0.1 (QIAGEN) with the following parameters: match score, 1; mismatch cost, 2; indel cost, 3; length fraction, 0.8; and similarity fraction, 0.9. The number of raw read pairs per sample and the ratio of reads mapped on the reference sequences were shown in Figure 1B and Supplementary Table S2. Non-specific reads mapped to multiple locations were ignored for the analysis presented in Figure 1B and Supplementary Table S2, while all reads were used in Figure 1CD and Supplementary Figure S1. Original sequence reads were deposited in the DRA/SRA database with the following accession numbers (Library A: DRS268643, Library B: DRS268645, and Library C: DRS268650). The accession number for the BioProject was PRJDB11466.

**Figure 1.**
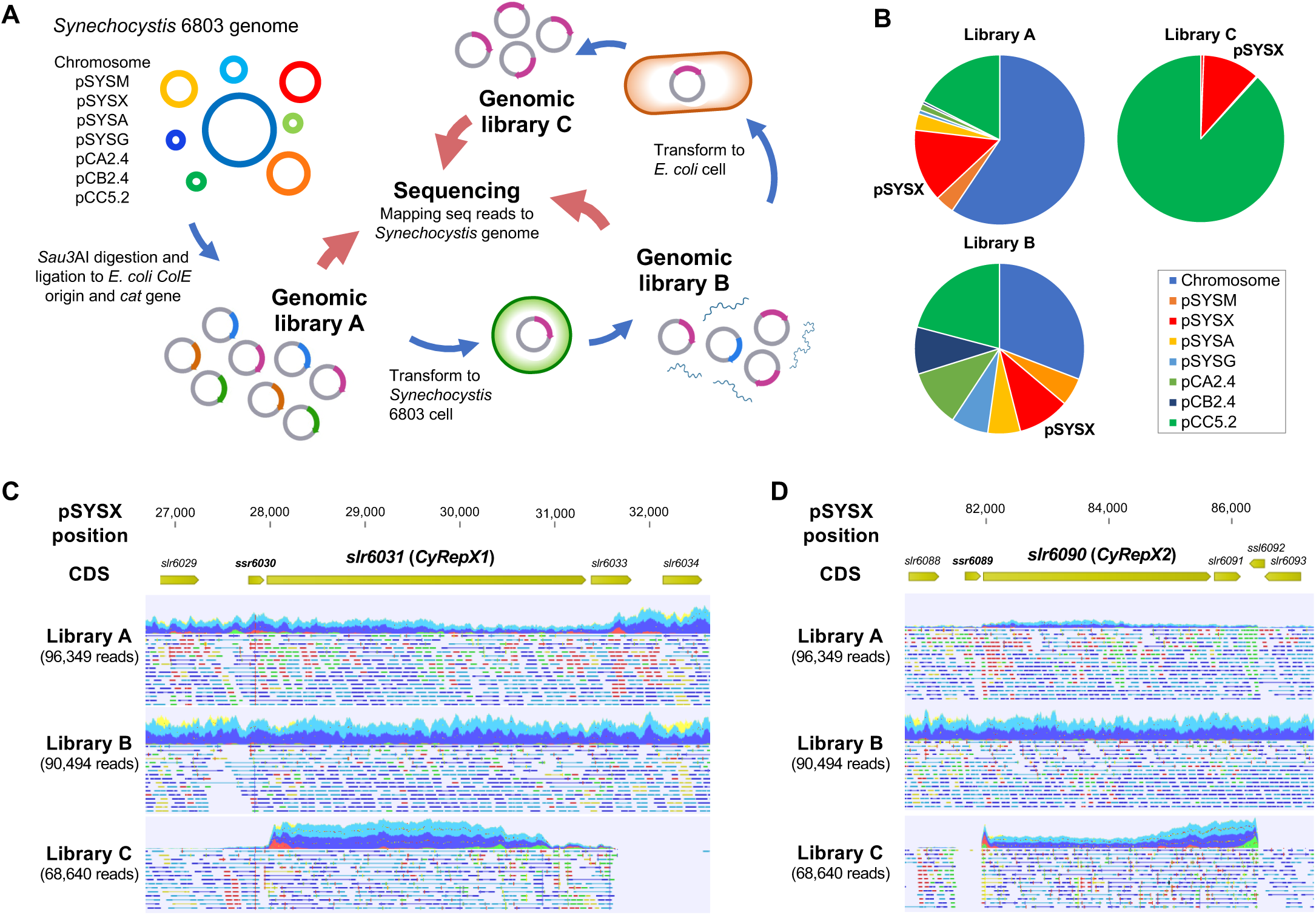
Screening of autonomous replication region in *Synechocystis* 6803 cell. (A) Scheme of autonomous-replication sequencing (AR-seq). Library A, prepared from restriction enzyme-digested *Synechocystis* 6803 genomic DNA, was transformed into *Synechocystis* 6803 cells, and library B was obtained from the resulting transformants. Library B was transformed again into *E. coli* to obtain library C containing clones with autonomous replication activity. Libraries A-C were subjected to quantitative sequence analysis, and the resulting sequence reads were mapped to the *Synechocystis* 6803 genome sequence as a reference. (B) The composition of sequence reads of the *Synechocystis* 6803 genomic libraries obtained from AR-seq analysis. Three libraries were sequenced by MiSeq and the reads were mapped to the *Synechocystis* 6803 genome (chromosome and 7 plasmids, pSYSM, pSYSX, pSYSA, pSYSG, pCA2.4, pCB2.4, and pCC5.2). Library A: *S*. 6803 genomic library before screening, Library B: DNA extracted from *Synechocystis* 6803 cells transformed with Library A, Library C: DNA extracted from *E. coli* cells transformed with Library B. (C, D) Mapping results of sequence reads to pSYSX as references. pSYSX regions around *slr6031* (C) and *slr6090* (D) are shown. Reads that have been successfully mapped to pSYSX as a pair are shown as blue or light blue, while if only one of the reads in the pair has been mapped, it is shown as red or green.

### 2.3 Phylogenetic analysis

The amino acid sequences of Slr6031 and Slr6090 (accession numbers: BAD02088.1 and BAD02147.1) were obtained from the database, and the top 100 most similar sequences for each protein were obtained by NCBI BLAST search as candidate homologs. After excluding overlapping sequences, a total of 148 homologous sequences of Slr6031 and Slr6090 were obtained. Phylogenetic analysis was performed on these homologs together with 59 previously determined Rep proteins including CyRepA1 and CyRepA2 (Sakamaki et al. 2023a), the *Synechococcus* 7942 pUH24/pANS Rep and RSF1010 RepC, using ClustalW within MEGA11 with default parameters (Tamura et al. 2021). Evolutionary history was inferred using the neighbor-joining method (Saitou and Nei 1987). The optimal tree depicted in Figure 2A was drawn to scale, with branch lengths in the same units as those of the evolutionary distances used to infer the phylogenetic tree. The evolutionary distances were computed using the Poisson correction method and expressed as the number of amino acid substitutions per site. This analysis involved 211 amino acid sequences (Slr6031, Slr6090 and their 148 homologs, 59 homologs of CyRepA1/2, pUH24/pANS Rep and RSF1010 RepC). All ambiguous positions were removed for each sequence pair (pairwise deletion).

**Figure 2.**
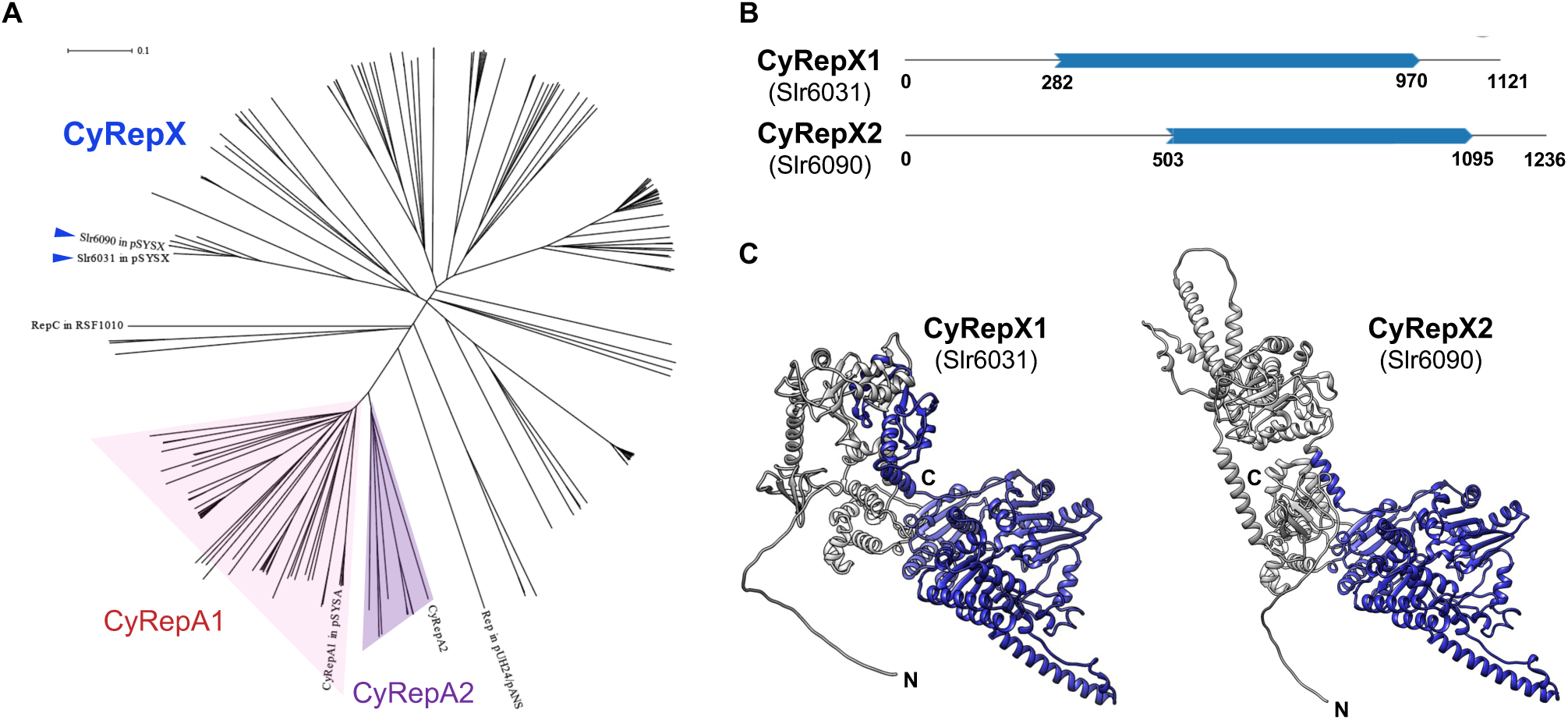
Phylogenetic and structural analysis of CyRepX1 (Slr6031) and CyRepX2 (Slr6090) (A) Phylogenetic tree of CyRepX. CyRepX1 (Slr6031), CyRepX2 (Slr6090) and its 148 homologs were used for phylogenetic analysis along with 59 CyRepA homologs, pUH24/pANS Rep, and RSF1010 RepC used as outgroups. The unrooted tree is drawn to scale, with branch lengths in the same units as those of the evolutionary distances used to infer the phylogenetic tree. CyRepX1 (Slr6031) and CyRepX2 (Slr6090) classified into same clade are indicated by arrow heads. (B) Structurally homologous region to the Rep-related CyRepX proteins predicted by Foldseek (van Kempen et al. 2023). CyRepX1 and CyRepX2 show structural similarity to A0A132Z0X2 and Q58352 in AFDB-PROTEOME, respectively, and they contain the DEXDc and HELICc motifs characteristic of Rep proteins. Blue arrows indicate the homologous region of CyRepX with Rep related proteins (A0A132Z0X2 and Q58352) shown in Supplemental Figure S2. The numbers indicate the position of the amino acid sequence in each protein. (C) 3D structures of CyRepX1 and CyRepX2 predicted using AlphaFold2. The 3D structures are blue color-coded based on the results of structural homology predictions by Foldseek.

### 2.4 Prediction of 3D structure of CyRepXs

AlphaFold2 was used for 3D structure prediction (AlQuraishi 2019, Cramer 2021). The predicted 3D structure of Slr6031 (CyRepX1), Slr6090 (CyRepX2), and the structural homologs (A0A132Z0X2 and Q58352) were obtained from the AlphaFold Protein Structure Database. Domain predictions were performed using SMART software (Letunic et al. 2021). Domains within the predicted 3D structures were colored using UCSF Chimera (University of California San Francisco, San Francisco, CA, USA) (Pettersen et al. 2004). Predicted 3D structure data of CyRepX1, CyRepX2 and the structural homologs (A0A132Z0X2 and Q58352) are provided in Supplementary Data 2-5.

### 2.5 Plasmid and strain construction

To construct the expression vectors p6031-1C-GFP and p6090-1C-GFP, a DNA fragment containing the *lacI* gene, the *trc* promoter, the *gfp ^mut2^* gene, the chloramphenicol resistance marker gene (*cat*), and the ColE origin, was PCR amplified using primer set F1/R2 (Supplementary Table S2) and the pYS1C-GFP plasmid as a template. The pSYSX region including either *slr6031-slr6032-slr6033* or *slr6090- slr6091* genes (27,879-32,722 and 82,597-87,154 in pSYSX: AP006585), which were suggested by AR-seq analysis to have autonomous replication activity, were also PCR amplified using appropriate primer sets (F3/F4 and F5/F6) and combined with the vector backbone by In-Fusion cloning. The vector sequences of p6031-1C-GFP and p6090-1C-GFP are available from Supplementary Data 6 and 7.

To construct the plasmid pYS4ST-GFP, which can be transferred by conjugation, the *oriT* region and pYS4S-GFP were PCR amplified using appropriate primers (F7/F8 and F9/F10) and templates (pUB307IP and pYS4S-GFP) (Itaya et al. 2018, Sakamaki et al. 2023a) and combined by In-Fusion cloning. Using the plasmid pYS4ST-GFP, the *oriT*-*Sp^R^*-*ColE* region was amplified using primer set F3/R11 and combined with the DNA fragment containing the *slr6090* region amplified using primer set F5/R6. The resulting plasmid p6090-ST was used for the compatibility test with p6031-1C-GFP. The sequence of p6090-ST is available from Supplementary Data 8.

### 2.6 Flow cytometry

Cells were analyzed by GFP fluorescence-activated cell sorting (FACS) using a BD Accuri^TM^ C6 Flow cytometer (BD Biosciences, San Jose, CA, USA) and BD CFlow software (BD Biosciences), as described by Watanabe et al. (2012) (Watanabe et al. 2012). Cyanobacterial cells showing chlorophyll fluorescence were sorted using the FL3 channel and GFP fluorescence was measured using the FL1 channel.

### 2.7 Microscopy

Fluorescence images were obtained using a BX53 microscope (OLYMPUS, Tokyo, Japan) at 100× magnification with a DP71 digital camera (OLYMPUS) and DP Controller software ver. 3. 3. 1. 292 (OLYMPUS).

### 2.8 DNA extraction

Genomic DNA including chromosome and plasmids was prepared from *Synechococcus* 7942 and *Synechocystis* 6803 according to a previous report (Itaya et al. 2003). We collected 25 mL of cyanobacteria culture by centrifugation (1,870 × *g*, 10 min) and resuspended the cell pellet in 4 mL of NaCl/EDTA solution (120 mM/ 50 mM, pH 8.0) and 1 mL of a saturated sodium iodide solution. After incubation for 30 min at 37 °C, the cells were washed twice with 5 mL of H2O and resuspended in 1 mL of NaCl/EDTA solution containing 10 mg/mL of lysozyme and 0.1 mg/mL RNase A final concentration, followed by incubation for 1 h at 37 °C. After an overnight incubation with 10 µL of proteinase K at 20 mg/mL and 1.0% sodium dodecyl sulfate (SDS), an equal volume of a phenol/chloroform mixture was added and mixed well, followed by centrifugation (10,000 × *g*, 10 min). The upper phase was again treated with phenol/chloroform mixture, transferred to a fresh tube, and an equal amount of 2-propanol with 0.3 M sodium acetate was then added to precipitate the genomic DNA. DNA was collected by centrifugation (10,000 × *g*, 10 min), rinsed with 70% ethanol, and air dried. The DNA pellet was dissolved in TE buffer and used for the following analyses.

### 2.9 Estimation of plasmid/chromosome ratios

To compare the autonomous replication activity of CyRepA and CyRepX proteins, the ratio of plasmid to chromosome copy number in *Synechocystis* 6803 or *Synechococcus* 7942 cells was estimated by droplet digital PCR using the QX200 system (Bio-Rad). Primer sets (F12/F13 and F14/R15, Supplementary Table S1) and HEX-conjugated fluorescent oligo probes (oligo 1 and oligo 2) targeting the *sigA*/*rpoD1* gene region (*sll0306* and *Synpcc7942_1557*) were used for the estimation of chromosome numbers in *Synechocystis* 6803 or *Synechococcus* 7942. To investigate the copy number of plasmids, pYS1C-GFP, p6031-1C-GFP, and p6090-1C-GFP, primer set F16/R17 and FAM-conjugated oligo probe (oligo 3) targeting the *cat* gene region were used. For the VIII23 plasmid, primer set F18/R19 and FAM conjugated oligo probe 4 targeting the *aph* gene were used. The TaqMan PCR reaction mixture was assembled from a 2× ddPCR master mix (Bio-Rad), 20× primers and probes (final concentrations of 900 and 250 nM, respectively) and template (variable volume) in a final volume of 20 μL. Each assembled ddPCR reaction mixture was then loaded into the sample well of an eight-channel disposable droplet generator cartridge (Bio-Rad). A volume of 70 μL of droplet generation oil (Bio-Rad) was loaded into the oil well for each channel. The cartridge was placed into the droplet generator (Bio-Rad). The cartridge was removed from the droplet generator, where the droplets that collected in the droplet well were then manually transferred with a multichannel pipet to a 96-well PCR plate. The plate was heat sealed with a foil seal (Bio-Rad) and then placed on a conventional thermal cycler and amplified to the end point (40 cycles). After PCR, the 96-well PCR plate was loaded on the droplet reader (Bio-Rad), which automatically reads the droplets from each well of the plate (32 wells/h). Analysis of the ddPCR data was performed with QuantaSoft analysis software (Bio-Rad) that accompanied the droplet reader.

The estimation of chromosomal copy numbers was performed as previously described (Gartner et al. 2019, Klotz et al. 2016, Watanabe and Yoshikawa 2016). The cyanobacterial cells were fixed with 0.005% Tween 20 and 1% glutaraldehyde for 30 min on ice. After washing with 1 mL of PBS, cell pellets were frozen and then thawed at 25 °C. The cell pellets were resuspended in 50 µL of 10 µM SYTOX Green solution, which was diluted with 50 mM trisodium citrate (pH 8.0). After incubation for 12 h at 4 °C, the cells were subjected to flow cytometry. *Synechococcus* 7942 cells cultured in phosphate-depleted BG-11 medium for one week with one chromosome copy per cell were used as standards (Watanabe et al. 2015, Watanabe and Yoshikawa 2016).

## 3 Results

### 3.1 Library screening of sequences responsible for autonomous replication activity in *Synechocystis* 6803

To gain new insights into autonomously replicating regions in the *Synechocystis* 6803 genome, we conducted Autonomous Replication sequencing (AR-seq), which combines library screening and sequencing (Figure 1A) (Sakamaki et al. 2023a). *Synechocystis* 6803 genomic library (library A) containing 2.5 × 10^4^ clones was transformed to *Synechocystis* 6803 cells, and 6.3 × 10^3^ of independent clones were obtained. DNA extracted from *Synechocystis* 6803 transformants was pooled as library B, which contained the autonomously replicating sequences in addition to *Synechocystis* 6803 genomic DNA. To isolate plasmids that replicated independently of chromosomal integration in *Synechocystis* 6803 cells, library B was introduced into *E. coli*, and only the plasmid DNA was pooled as library C.

Next, a comprehensive sequencing analysis was performed to reveal the genomic regions in the libraries. The results showed that pSYSX occupied a major proportion of the library in the *Synechocystis* 6803 genomic library A (Figure 1B and Supplementary Table S2), suggesting a high copy number of pSYSX in our *Synechocystis* 6803 strain among the large plasmids, as previously reported (Sakamaki et al. 2023a). The plasmid copy number in library B was more evenly spread than in library A, while in library C, the majority of sequence reads were assigned to pSYSX (10.97%) and pCC5.2 (88.19%) (Figure 1B and Supplementary Table S2) (Jin et al. 2018, Sakamaki et al. 2023a). Because it was already known that pCC5.2 contained a region with high replication activity (Sakamaki et al. 2023a), the following experiments were focused on pSYSX. We mapped the sequence reads to pSYSX to identify the region required for replication and observed that two regions containing full-length sequences of the *slr6031* and of the *slr6090-slr6091* genes were present in the sequence reads from libraries C (Figure 1C and D). The plasmid pSYSX contains a complete duplication (positions 4,092-27,731 and 58,810–82,449 bp) in a tandem arrangement (Kaneko et al. 2003), and the regions found by AR-seq were located adjacent to these duplicated regions (Supplementary Figure S1). While not identical, the Slr6031 and Slr6090 proteins showed high similarity to each other; however, their functions have not yet been specified. Thus, we conducted conservation and functional analysis of these proteins.

### 3.2 Phylogenetic analysis of Slr6031 (CyRepX1) and Slr6090 (CyRepX2) in pSYSX

To characterize Slr6031 and Slr6090 in pSYSX, we performed a sequence-based analysis of the conserved regions and structure of these proteins. Proteins showing homology to Slr6031 and Slr6090 obtained from BLAST analysis were subjected to phylogenetic analysis along with CyRepA1 and CyRepA2 and other Reps identified in cyanobacteria. Homologs of Slr6031 and Slr6090 were identified in various cyanobacteria, including *Synechococcales*, *Oscillatoriophycideae*, *Nostocales*, *Pleurocapsales*, *Pseudanabaenales*, *Spirulinales*, and *Pleurocapsales*, albeit with low sequence identity and placed in several clades distinct from that of CyRepA proteins (Figure 2A). These findings suggest that the protein family, including Slr6031 and Slr6090 has evolved into a more diverse set of proteins than CyRepA in cyanobacteria. Since analyses using SMART or Pfam databases did not identify specific domains for Slr6031 and Slr6090, the Foldseek algorithm was used to explore the structural homologs of these proteins (van Kempen et al. 2023). The results of searching the AFDB-Proteome database with Foldseek showed that Slr6031 and Slr6090 have structural homology to proteins in *Enterococcus faecium* (UniProt ID: A0A132Z0X2) and *Methanococcus jannaschii* (Q58352), respectively (Figure 2B). Interestingly, the regions in these proteins that showed similarity to Slr6031 and Slr6090 contain DEXDc and HELICc motifs, characteristic of helicase family proteins (Figure 2C and Supplemental Figure S2). Since the DEXDc domain is also a characteristic domain of the CyRepA proteins, which have autonomous replication activity in cyanobacteria (Sakamaki et al. 2023a), we expected that Slr6031 and Slr6090 would also act as Rep.

### 3.3 Construction and evaluation of autonomously replicating plasmid

To evaluate the replication activity of Slr6031 and Slr6090 in cyanobacteria, the expression vectors p6031- and p6090-1C-GFP (Supplementary Figure S3) were constructed based on a plasmid pYS1C-GFP containing the *E. coli* ColE1 origin of replication, the *cat* gene, and *gfp* gene under the control of the *trc* promoter with the *lacI* repressor gene (Sakamaki et al. 2023a). Each of these plasmids was introduced into *Synechococcus* 7942, and the obtained transformants were used for following assays. After 48 h of pre-incubation, the cells were cultivated for additional 2 h in presence of 1 mM IPTG, and subsequently subjected to fluorescence microscopy. We observed GFP fluorescence in *Synechococcus* 7942 cells carrying p6031- and p6090-1C-GFP as well as pYS1C-GFP (Figure 3A). Compared to pYS1C-GFP, GFP fluorescence of the p6031- and p6090-1C-GFP transformants were not uniform and some cells did not show fluorescence (Figure 3A, white arrows). The expression levels of GFP were estimated and compared by flow cytometry. Consistent with the microscopy results, GFP fluorescence of p6031- and p6090-1C-GFP transformants analyzed by flow cytometry was lower than that of pYS1C-GFP and contained cells without fluorescence (Figure 3B). To confirm the structure of p6031- and p6090-1C-GFP plasmids in *Synechococcus* 7942 cells, DNA was prepared from cells harboring plasmids and introduced into *E. coli*. Plasmids were extracted from the resulting *E. coli* transformants and compared with the initial p6031- and p6090-1C-GFP plasmids using restriction enzyme digestion (Figure 3C). The results indicated that both plasmids were identical (Figure 3C), suggesting that p6031- and p6090-1C-GFP were maintained as intact plasmids, and that the regions containing *slr6031* or *slr6090* conferred autonomous replication activity in *Synechococcus* 7942.

**Figure 3.**
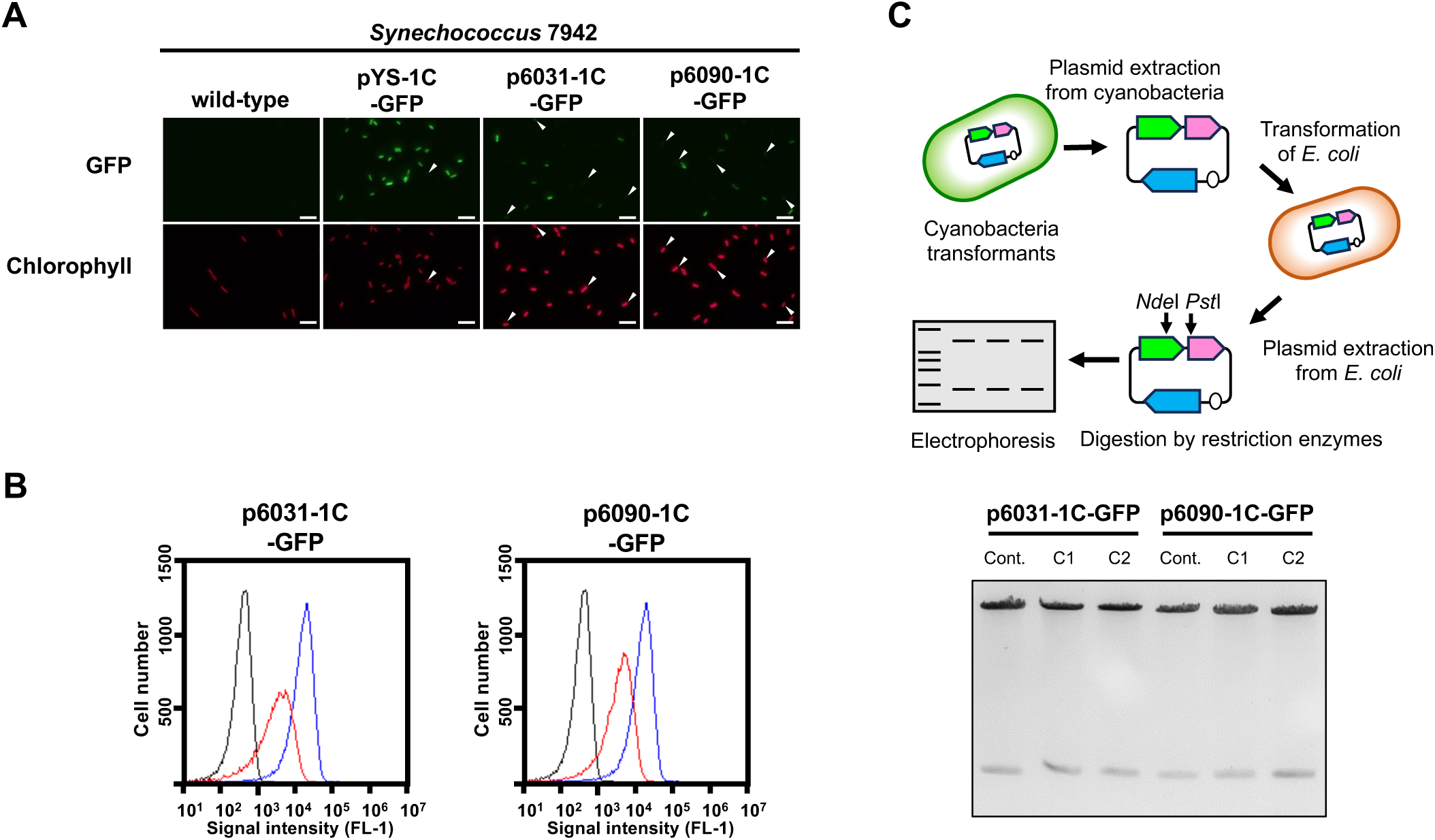
Autonomous replication activity of CyRepX in heterologous cyanobacterial host *Synechococcus* 7942 cells. *Synechococcus* 7942 cells harboring pYS1C-GFP, p6031-1C-GFP and p6090-1C-GFP were cultivated for 2 days. The cells were harvested 2 hours after addition of 1 mM IPTG and compared the GFP and chlorophyll with wild-type strain. (A) Fluorescence microscopy images. White bar:10 μm (B) FACS analysis of GFP fluorescence. Signal intensity of FL1 indicating GFP fluorescence in *Synechococcus* 7942 wild-type (black) pYS1C-GFP (blue) and p6031-1C-GFP or p6090-1C-GFP transformants (red) cultivating with IPTG are shown. (C) Verification of plasmid structure maintained in cyanobacterial cells. Electrophoresis image of plasmids digested by the restriction enzymes is shown with the scheme of the analysis of plasmid structure. To determine whether plasmids are maintained in cyanobacterial cells in a circular structure, DNA was extracted from cyanobacteria transformants carrying plasmids (p6031-1C-GFP and p6090-1C-GFP) and introduced into *E. coli*. After the plasmid was extracted from the *E. coli* cells, plasmids were digested with restriction enzymes *Nde*I and *Pst*I, and compared to the plasmids before the transformation of cyanobacteria. The plasmids before transformation were used as controls. The results of two independent clones are shown as C1 and C2.

The possible maintenance of p6031- and p6090-1C-GFP plasmids in *Synechocystis* 6803 was evaluated using the same approaches as in *Synechococcus* 7942. Although *Synechocystis* transformants of p6031- and p6090-1C-GFP exhibiting GFP fluorescence were obtained (Figure S4A and B), no colonies were observed when the DNA extracted from the p6031-1C-GFP transformant was introduced into *E. coli*. In the case of p6090-1C-GFP, only two colonies were obtained, one of which had a distorted plasmid structure (Figure S4C). These results indicated that the majority of p6031- and p6090-1C-GFP constructs were not maintained as plasmids but likely integrated into the endogenous *slr6031* or *slr6090* loci. Since *Synechococcus* 7942 replicated the plasmids as intact plasmids, it was used as host for most subsequent analyses.

### 3.4 Estimation of plasmid copy number

Next, we estimated the copy number of our plasmids by droplet digital PCR technology. The *sigA/rpoD1* gene encoding the RNA polymerase sigma factor was used to estimate chromosome copy number, while the *cat*, Cm resistance marker gene, encoded on each plasmid was utilized to determine plasmid number. In *Synechococcus* 7942, the copy number of pYS1C-GFP with CyRepA2 was 34.1 times higher than that of the chromosome, whereas those of p6031-1C-GFP and p6090-1C-GFP carrying *slr6031*/*slr6090* regions were 4.1 and 6.3 times higher, respectively (Table 1). The chromosome copy number of *Synechococcus* 7942 under this condition was 2-6 copies per cell (Supplementary Figure S5), suggesting that the copy number of pYS1C-GFP, p6031-1C-GFP and p6090-1C-GFP in *Synechococcus* 7942 cells were estimated as 68-204, 8-24, and 12-36 copies, respectively (Table 1). Similar experiments were performed in *Synechocystis* 6803 and showed that the copy number of pYS1C-GFP was 26-87 copies per cell (8.7 times as many copies as the chromosome with 3-10 copies per cell) (Table 1 and Supplementary Figure S5). We did not test copy numbers of the p6031- and p6090-1C-GFP constructs in *Synechocystis* 6803 because they were likely not maintained as individual plasmids (see preceding section 3.3). Furthermore, we estimated the copy number of the VIII23 plasmid, which possesses CyRepA1 and showed autonomous replication activity in both cyanobacteria (Kaltenbrunner et al. 2023). The ratio of chromosomes to VIII23 plasmid was examined by ddPCR targeting the *aph*, Km resistance marker gene, and it was found that in *Synechocystis* 6803, the plasmid copy number was similar to that of the chromosomes (Table 1), compared to 33-99 copies in *Synechococcus* 7942. Hence, the plasmid VIII23 with CyRepA1 existed at a lower copy number than plasmid pYS with CyRepA2 in both *Synechococcus* 7942 and *Synechocystis* 6803. These observations are consistent with the copy number of pSYSA (harboring CyRepA1) and pCC5.2 (harboring CyRepA2) in *Synechocystis* 6803 cells (Nagy et al. 2021), although direct comparisons may not be appropriate due to the different plasmid configurations.

**Table 1.** Copy number of chromosome and plasmid in transformants.

|  | Chromosome<br>copy number | Ratio<br>(plasmid per chromosome) | Estimated plasmid<br>copy number |
| --- | --- | --- | --- |
| *Synechococcus* 7942<br>pYS1C-GFP | 2-6 | $34.1 \pm 1.6$ | 68-204 |
| *Synechococcus* 7942<br>p6031-1C-GFP | 2-6 | $4.1 \pm 0.2$ | 8-24 |
| *Synechococcus* 7942<br>p6090-1C-GFP | 2-6 | $6.3 \pm 0.7$ | 12-36 |
| *Synechococcus* 7942<br>VIII23 | 2-6 | $16.5 \pm 0.4$ | 33-99 |
| *Synechocystis* 6803<br>pYS1C-GFP | 3-10 | $8.7 \pm 0.2$ | 26-87 |
| *Synechocystis* 6803<br>VIII23 | 3-10 | $0.8 \pm 0.04$ | 2-8 |

The experiments were performed using DNA extracted from cells after 2 days of incubation (O.D.<sub>750</sub> = 0.6-1.0) in BG11 liquid medium containing antibiotics (Cm or Km). Chromosome abundance was quantified by ddPCR targeting the *sigA/rpoD1* gene on *Synechococcus* 7942 and *Synechocystis* 6803 chromosomes, while plasmid abundance was determined by targeting the drug resistance gene (*cat/aph* gene) carried on the respective plasmids, and the ratio (plasmid/chromosome) was calculated.

### 3.5 Comparison of plasmid maintenance in cyanobacterial cell

Next, the stability of the plasmid was examined in individual cells with GFP fluorescence as a reporter. Due to the expected loss of plasmid in the medium lacking antibiotics for plasmid selection, *Synechococcus* 7942 cells carrying pYS1C-GFP, p6031-1C-GFP and p6090-1C-GFP were grown on the BG-11 plate containing IPTG with and without chloramphenicol (Cm) to estimate the GFP fluorescence. While GFP fluorescence was maintained under conditions with Cm, *Synechococcus* 7942 grown on Cm-free medium for three days reduced the share of cells showing fluorescence and increased the share of cells without fluorescence (Figure 4A). After an additional three days incubation under the Cm-free conditions, the population of no GFP fluorescence cells further increased, even in the cells harboring pYS1C-GFP (Figure 4B). In contrast to *Synechococcus* 7942, there was no reduction of GFP fluorescence in *Synechocystis* 6803 cells carrying pYS1C-GFP even after two consecutive transfers in Cm-free medium (Figure 4). pYS is originally derived from endogenous plasmid pCC5.2 in *Synechocystis* 6803, and it has been indicated that both pYS and pCC5.2 plasmids coexist in the cell (Sakamaki et al. 2023a). These observations imply that pYS has an active mechanism to avoid incompatibility with pCC5.2 and to be maintained in *Synechocystis* 6803 cells.

**Figure 4.**
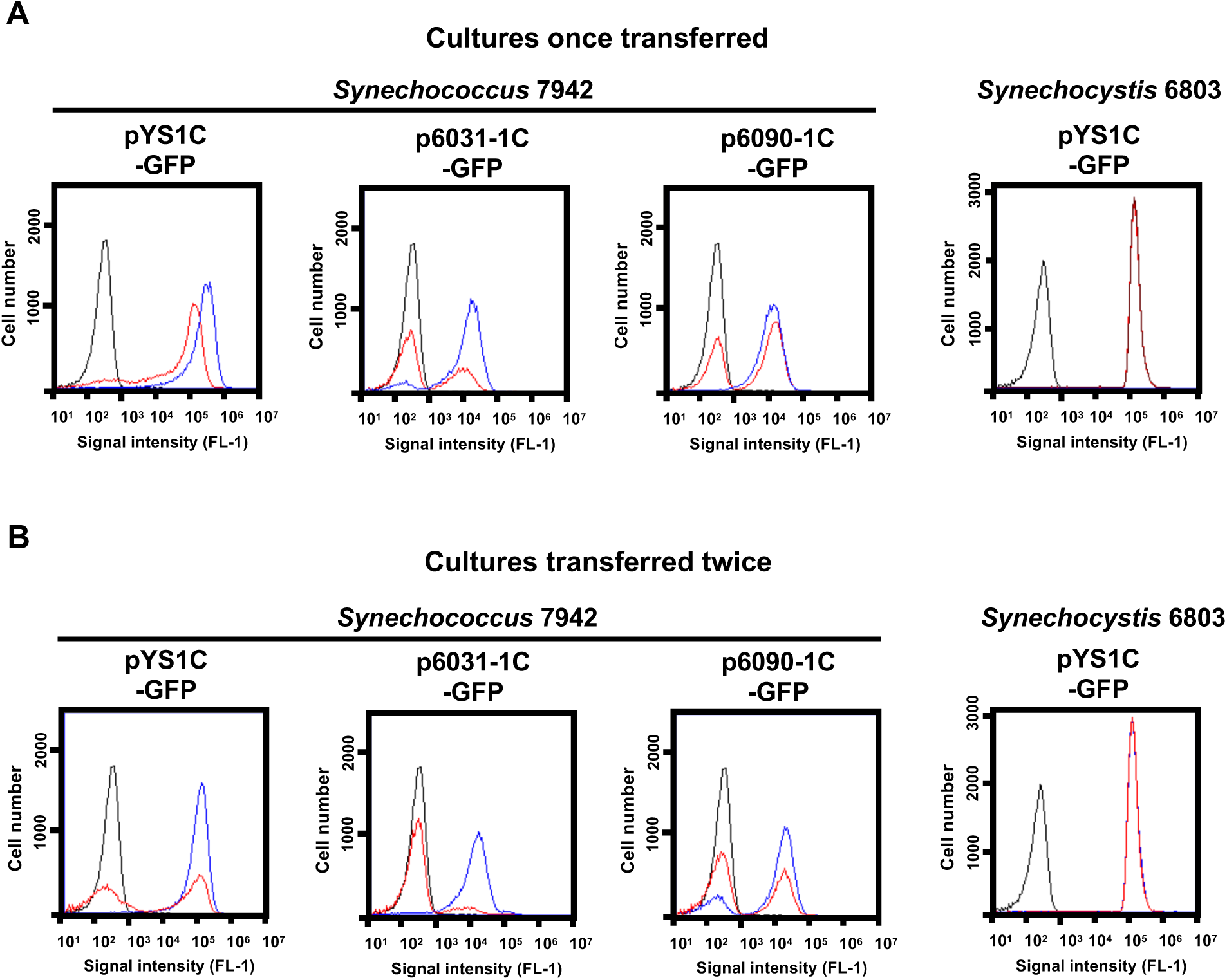
Comparison of plasmid maintenance in cyanobacterial cells. To compare the stability of the plasmids, each transformant was grown for 3 days on BG-11 plates containing 1 mM IPTG with (+) and without Cm (-) (Cultures once transferred). Transformants were again transferred to fresh BG11 plates with the same conditions and cultured for an additional 3 days (Cultures transferred twice). (A, B) FACS analysis of GFP fluorescence. Signal intensity of FL1 indicating GFP fluorescence in the plasmid transformants of *Synechococcus* 7942 and *Synechocystis* 6803 cultivated with (blue) and without Cm (red) are shown. The profiles drown in black indicates the FL1 fluorescence of the wild-type cells.

### 3.6 Compatibility of multiple plasmids in *Synechococcus* 7942

In *Synechocystis* 6803 cells, seven different plasmids coexist without showing incompatibility and there is likely little competition among plasmid Reps. We examined whether *Synechococcus* 7942 could simultaneously maintain multiple plasmids with different Reps. A VIII23 plasmid carrying *CyRepA1* and *aph* genes (Kaltenbrunner et al. 2023) was used to the compatibility test along with p6031-1C-GFP. Following the transformation of *Synechococcus* 7942 cells carrying p6031-1C-GFP with VIII23, we obtained colonies showing resistance to both Cm and Km, and confirmed that both plasmids are maintained in the correct structure (Figure 5A). Furthermore, additional introduction of the pEX2-mScarlet plasmid (Sakamaki et al. 2023a) harboring another Rep and Sp resistance marker gene *aad* into the cells harboring the p6031-1C-GFP and VIII23 plasmids resulted in transformants, and the three plasmids were maintained correctly (Figure 5B). These results indicate that *Synechococcus* 7942 is capable of carrying at least three plasmids.

**Figure 5.**
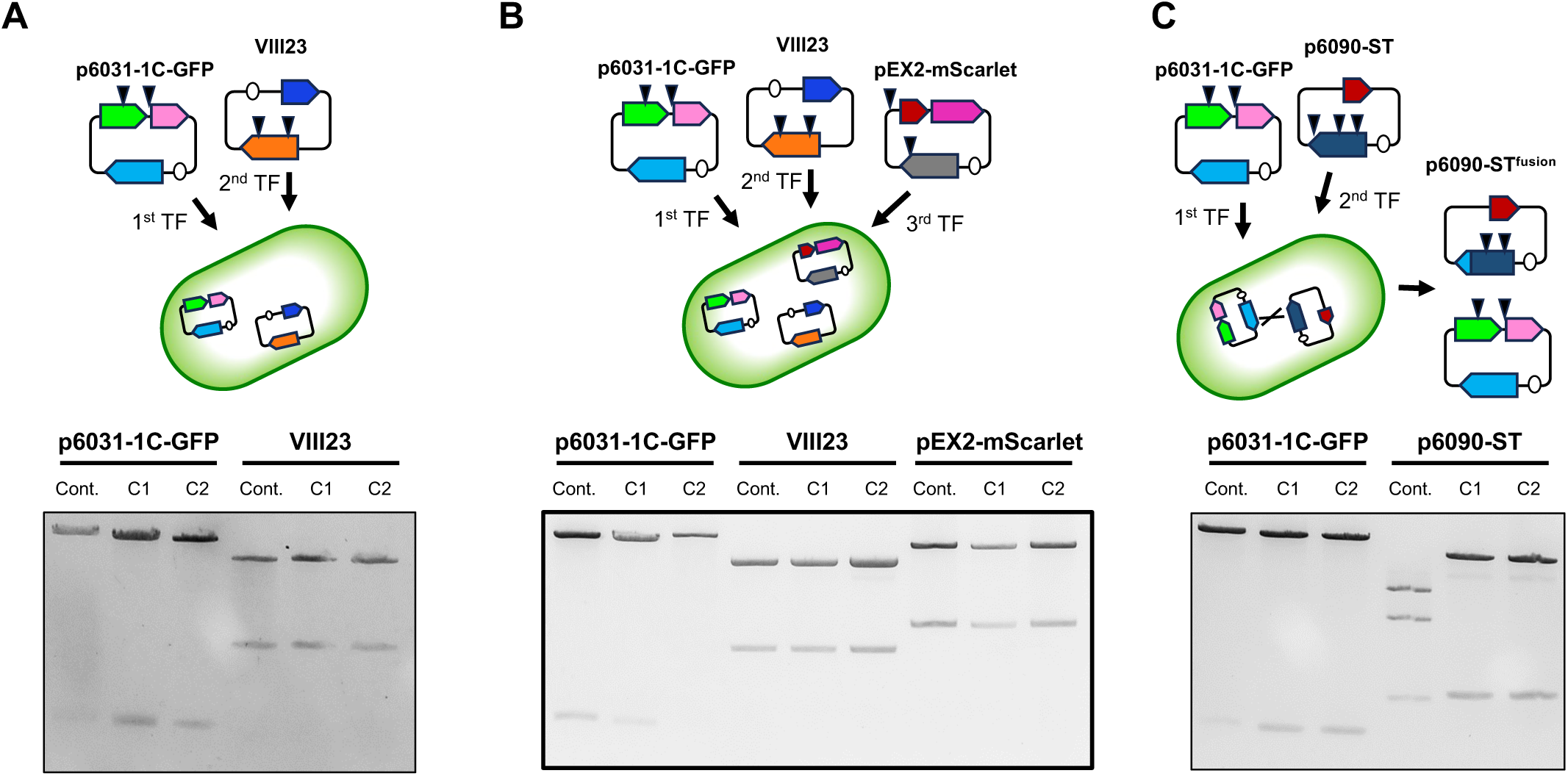
Multiple plasmid maintenance in *Synechococcus* 7942. Simultaneous maintenance of multiple plasmids, p6031-1C-GFP (containing *slr6031* and *cat*), VIII23 (*CyRepA1* and *aph*) (Kaltenbrunner et al. 2023), pEX2-mScarlet (pANS Rep gene and *aad*) (Sakamaki et al. 2023a) and p6090-ST (*slr6090* and *aad*), was studied in *Synechococcus* 7942. The plasmids prepared as described in Figure 3C were digested by appropriate restriction enzymes (p6031-1C-GFP: *Nde*I and *Pst*I; VIII23: *Xba*I; pEX2-mScarlet and p6090-ST: *Hind*III) and subjected to the electrophoresis. Arrow heads in the upper schemes indicate the locations recognized by each restriction enzyme. The results of two independent clones are shown as C1 and C2. (A, B) The simultaneous maintenance of p6031-1C-GFP, VIII23 and pEX2-mScarlet in *Synechococcus* 7942 cell. (C) The simultaneous maintenance of p6031-1C-GFP and p6090-ST. The electrophoresis pattern in p6090-ST changed, suggesting structural re-constitution of *slr6090* has been occurred.

We further tested whether two Rep candidates, Slr6031 and Slr6090, could be utilized simultaneously in *Synechococcus* 7942. The plasmid p6090-ST harboring the *slr6090* and *aad* genes was introduced into cells carrying p6031-1C-GFP, resulting in a strain resistant to both Cm and Sp. The plasmid structure of p6031-1C-GFP obtained from *Synechococcus* 7942 was comparable to that of the control. However, the structure of p6090-ST was altered. Sequence analysis showed that part of the *slr6090* region in p6090-ST (85,039-87,154 in pSYSX) was replaced by the homologous region of p6031-1C-GFP (30,391-32,722 in pSYSX) causing a difference in the *Hind*III cleavage pattern (Figure 5C). The resulting plasmid p6090-ST^fusion^ was likely generated by recombination (Supplementary Figure S3, Supplementary Data 9), forming a fusion protein of Slr6090 and Slr6031, which was possibly more suitable for *Synechococcus* 7942. In summary, two very similar plasmids, p6031-1C-GFP and p6090- ST^fusion^, resulting from microevolution, were shown to be maintained simultaneously.

## 4 Discussion

While numerous cyanobacteria maintain large plasmids, insight into the Rep proteins orchestrating the initiation of their replication has remained profoundly restricted. In this study, we performed AR-seq, a comprehensive genomic library screening approach, in *Synechocystis* 6803 and found two candidate Rep, Slr6031 and Slr6090, encoded in the pSYSX plasmid. Slr6031 and Slr6090 exhibit obvious autonomous replication activity and have distinct sequences and features from the known CyRepA1 and CyRepA2. Therefore, we propose to designate Slr6031 and Slr6090 as CyRepX1 (<u>Cy</u>anobacterial <u>Rep-</u>related protein encoding pSYS<u>X</u>) and CyRepX2, respectively. Intriguingly, despite the lower similarity among its homologs, CyRepX-type Rep proteins have maintained their presence across various genera of cyanobacteria, suggesting that their distinctly different evolution compared to the CyRepA family proteins (Figure 2A).

CyRepX1 and CyRepX2 exhibit substantial similarity with each other. Most remarkably, both are encoded within the large plasmid pSYSX, indicating the parallel presence of two very close replication initiation systems on a singular plasmid. The plasmid pSYSX manifests an intriguing structural peculiarity, with a tandem duplication encompassing more than a third of the plasmid (Supplementary Figure S1) (Kaneko et al. 2003). This duplication is distinctively unique to *Synechocystis* 6803. Even its closely related strain, *Synechocystis* sp. PCC 6714, bears a divergent gene composition in its large plasmids pSYLA, pSYLB and pSYLC (Kopf et al. 2014b) and it has a single CyRepX homolog (ORF ID: D082_40540) encoded on plasmid pSYLA (Supplementary Data 1). In contrast, in *Synechocystis* 6803, CyRepX1 and CyRepX2 are situated proximate to the duplicated region. Therefore, the dual CyRepX configuration with duplicated regions within pSYSX might have been acquired in a recent evolutionary process. Notably, within *Synechococcus* 7942, co-maintenance of p6031-1C-GFP and p6090-ST triggered a structural adjustment, primarily affecting plasmid p6090-ST, resulting in the fusion of Slr6090 and Slr6031 (Figure 5C and Supplementary Figure S3). Evidently, the interplay between host factors and the coexistence of other Reps potentially influenced Rep optimization.

The phenomenon of “multiple Reps in one plasmid” observed in pSYSX, is also found in clinically relevant bacteria, where it is associated with the spread of antimicrobial resistances. Especially the pSM19035 plasmid, belonging to the Inc18 family in *Firmicutes* and involved in erythromycin and lincomycin resistance, contains duplicated replication regions {Ceglowski, 1993 #768;Lioy, 2010 #767}). Although identical Reps have been observed in this plasmid, the precise physiological significance of these duplicated replication regions remains unclear. Similarly, plasmids of the IncF incompatibility group, which are prevalent in clinical enteric bacteria and contribute to the dissemination of antimicrobial resistance, also exhibit a unique feature (Villa et al. 2010). These plasmids have multiple replicons of different types, allowing them to replicate in a wide range of hosts. For instance, the IncF plasmid pGSH500 contains two functional replicons, one related to the narrow host range and the second to the broad host range (Osborn et al. 2000). This strategy of multiple replicons is important for expanding the plasmids’ host range and can lead to the replacement or the elimination of incompatible plasmids in certain situations. Plasmid pSYSX, characterized by a duplicated region and two copies of CyRepX, possibly arose by a similar principle as the Inc18 family plasmids. However, pSYSX is unique in its substantial size in comparison to Inc18 family plasmids. The tandem action of two Reps in pSYSX likely contributes to maintaining a relatively high copy number of the 106 kbp plasmid within *Synechocystis* 6803 (Figure 1B) (Nagy et al. 2021).

To reveal how frequent the arrangement with “multiple Reps in one plasmid” is in cyanobacteria, we investigated the conservation of CyRepA and CyRepX in 145 cyanobacteria with complete genome information available in the KEGG database. The number of proteins with full-length or partial domains of CyRepA1 (Slr7037) and CyRepX1 (Slr6031) and their respective ORF IDs are presented for each chromosome and plasmid (Supplementary Data 1). Within this dataset, comprising a total of 256 plasmids distributed across 145 cyanobacterial species, we found that 41 plasmids distributed within 26 cyanobacterial species belonging to the *Synechococcales*, *Pseudanabaenales*, *Oscillatoriophycideae*, *Nostocales*, and *Pleurocapsales*, possess multiple Reps in one plasmid. Of those, 33 plasmids had multiple CyRepA or CyRepX homologs, thereby indicating the prevalence of “multiple Reps in one plasmid” among the respective cyanobacteria. Notably, several large plasmids with multiple CyRepA and CyRepX were identified in *Acaryochloris marina* MBIC11017. Seven of the eight large plasmids in this cyanobacterium (except pREB6) harbor multiple genes for CyRepA and CyRepX homologs (Supplementary Data 1). Another interesting finding was that homologs of CyRepA and CyRepX proteins are widely distributed in cyanobacterial chromosomes as well as in plasmids (Supplementary Data 1). CyRepA homologs were identified on chromosomes in 80 of the 145 analyzed cyanobacteria. Eight of those cyanobacteria had a CyRepX homolog along with CyRepA, whereas no cyanobacteria were found to encode only CyRepX in their chromosomes. Further validation of the function of these chromosomally encoded CyRep homologs is needed.

Using AlphaFold2 and the Foldseek databases (van Kempen et al. 2023, Varadi et al. 2022), we identified a protein with structural similarity to CyRepX and predicted its function, indicating that structure-based prediction is useful for estimating the function of proteins of unknown function. CyRepX showed similarity to the helicase-related domains of its structural homologs, DEXDc and HELICc (UniProt ID: A0A132Z0X2 and Q58352, Supplemental Figure S2). Since CyRepA proteins also possess DEXDc domains (Sakamaki et al. 2023a), it expectedly found cyanobacterial Reps. Notably, the Rep proteins found in cyanobacteria do not show similarity to any of the Inc family plasmid proteins that have been studied and classified over the years, appearing to form a unique group (Shintani et al. 2015). Since there are several plasmids for which Reps have not yet been identified, further structure-based predictions coupled with experimental validation could reveal the diverse and unique set of Rep proteins in cyanobacteria.

The mechanism of CyRepX-mediated pSYSX replication initiation must be investigated in more detail in future studies. In case of pSYSA replication, previous studies have illuminated the role of two small RNAs, the *ssr7036* transcript and asRNA1. These elements, located upstream of the *CyRepA1* gene, are thought to be involved in regulating CyRepA1-mediated initiation of pSYSA replication, suggesting a θ-type replication mode similar to ColE (Kaltenbrunner et al. 2023). A model has been proposed that one transcript promotes pSYSA replication initiation, while the other RNA regulates it negatively. Interestingly, there are two pairs of very abundant sRNAs on pSYSX. Two of these originate in antisense orientation ∼25 nt downstream of the annotated start codons of *slr6031* (CyRepX1) and *slr6090* (CyRepX2) genes, while another two originate more distantly from *slr6031* and *slr6090* on the forward strand (Kopf et al. 2014a). The previous classification of transcripts assigned single transcriptional units (TUs) to each of these four transcripts, TU6027 and TU6028, as well as TU6082 and TU6083 (Kopf et al. 2014a). The respective TUs on forward and reverse strands have overlaps of 249 or 285 nt with each other (Figure 6A and B). Because they are located directly adjacent and partially overlap the *slr6031* and *slr6090* genes, it is likely that they play a role in the control of pSYSX replication initiation in *Synechocystis* 6803. The anti-Shine-Dalgarno sequences and also the segments complementary to the start codons are both in a much less structured region of these asRNAs than the adjacent stem-loop structures on both sides (Figure 6C). This indicates that they can likely base pair with the mRNAs in these regions critical for the initiation of translation and hence regulate expression of CyRepX1 and CyRepX2. More empirical testing is needed to further substantiate this hypothesis.

**Figure 6.**
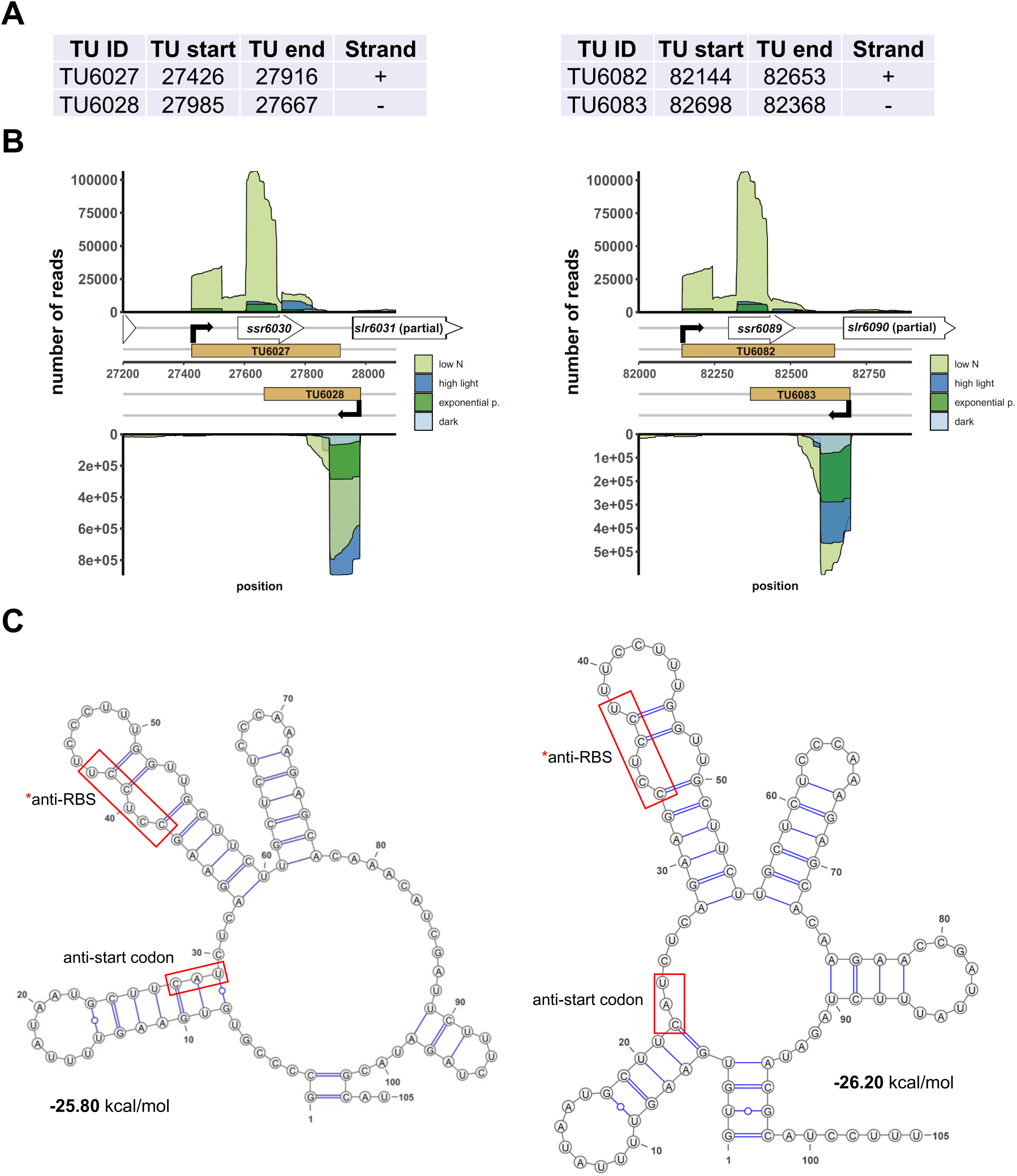
Two abundantly transcribed loci with overlapping transcripts on plasmid pSYSX in *Synechocystis* 6803. (A) Definition of the transcriptional units TU6027, TU6028, as well as TU6082 and TU6083 according to previous analyses by dRNA-seq (Kopf et al. 2014a). The TU start and end positions are given for the *Synechocystis* 6803 pSYSX sequence available in GenBank under accession number AP006585. (B) Location of TUs (indicated by light brown rectangles) according to the previous annotation of the transcriptome and genome-wide mapping of transcriptional start sites (Kopf et al. 2014a), together with the mapping data from previous dRNA-seq analyses for four conditions, nitrogen starvation, low N (light green), high light (470 µmol photons m^-2^s^-1^ for 30 min, blue), exponential growth phase (green) and 12 h darkness (light blue). Transcriptional start sites at the beginning of the TUs are indicated by bent black arrows. Note that TU6027 and TU6082 belong to the duplicated plasmid segment and therefore are identical in sequence. (C) The secondary structures and minimum free energy values in kcal/mol were calculated for the first 105 nt of the two asRNAs by RNAfold (Gruber et al. 2015) with default parameters and visualized with VARNA version 3.93 (Darty et al. 2009). The segments matching the likely Shine-Dalgarno sequences (anti-RBS) and start codons of *slr6031* and *slr6090*, respectively, are boxed in red.

The study of cyanobacterial plasmid replication will facilitate their application. Cyanobacteria, convert CO2 into organic molecules through photosynthesis, offering them promise for the carbon-neutral production of diverse compounds. In this study, we obtained also valuable insights into the utilization of vectors constructed with Reps encoded by three plasmids of *Synechocystis* 6803. First, the copy number was estimated in the plasmid vectors carrying CyRepA or CyRepX. As expected, the pYS vector with CyRepA2 had the highest copy number in both *Synechococcus* 7942 and *Synechocystis* 6803 (Table 1). This is consistent with the expression levels examined using GFP, again indicating that pYS is a suitable vector plasmid for gene overexpression (Sakamaki et al. 2023a). The copy number of VIII23 with CyRepA1 is comparable to the chromosome copy number in *Synechocystis* 6803, while it was found to have a copy number 10-fold higher than the chromosome in *Synechococcus* 7942 (Table 1). Since the VIII23 plasmid contains a regulatory region that expresses an sRNA along with CyRepA1, it is necessary to compare the copy numbers of pYS and VIII23 in a same construct to compare the function and activity between CyRepA1 and CyRepA2. The CyRepX plasmids, p6031-1C-GFP and p6090-1C-GFP were found to have a 4- and 6-fold higher copy number than the chromosome in *Synechococcus* 7942 (Table 1). These plasmids clearly have a lower copy number than pYS; thus, they can be used for different purposes, such as fine-tuning the expression levels of target genes.

Second, we addressed the differences in the stability of plasmids with CyReps (Figure 4). In *Synechococcus* 7942, the expression level of GFP used as a reporter gradually decreased in the medium without Cm used as a selection marker. GFP fluorescence was lost earlier for p6031 and p6090 plasmid vectors with low copy number than for pYS with high copy number. Consistent with this observation, GFP fluorescence in *Synechococcus* 7942 cells carrying the p6031 and p6090 plasmids was inconsistent (Figure 3AB), suggesting that these plasmids are unstable compared to pYS. This instability is a potentially useful property for transient expression of cytotoxic proteins (e.g., Cas9 used for genome editing). Interestingly, *Synechocystis* 6803 carrying pYS plasmid showed minimal reduction in GFP fluorescence when cultured in Cm-free medium, implying the existence of an unknown mechanism that stably maintains this plasmid. Our previous results showed that pYS was maintained along with original plasmid pCC5.2 in *Synechocystis* 6803 cells (Sakamaki et al. 2023a). pYS contains additional genetic parts, such as an *E. coli ColE* origin and a *gfp* expression system. These elements may affect the stability of pYS in *Synechocystis* 6803 cells. Identifying reasons for stable maintenance of pYS will contribute to vector development in *Synechocystis* 6803 and other cyanobacteria.

Finally, we obtained information on the simultaneous maintenance of multiple plasmids with different Reps (Figure 5). In *Synechococcus* 7942, CyRepX1 and CyRepX2, CyRepA1, and a pEX2 plasmid with a Rep derived from the endogenous pANS could be maintained together (Figure 5A and 5B). The plasmids with CyRepX1 and CyRepX2 were also successfully maintained simultaneously, although unexpectedly a recombination was observed in CyRepX2 (Figure 5C and Supplementary Figure S3). These results showed that *Synechococcus* 7942 allows multiple plasmids without causing incompatibility. Hence, the combination of multiple, different vectors becomes available, allowing for more complex genetic modifications and synthetic biology applications.

## 5 Data availability statement

The data presented in the study are deposited in the DDBJ repository, accession number PRJDB11466.

## 7 Author contributions

SW: design of this study. OK, MS, YS KM and KN-M: data curation, experiment, and writing— original draft preparation. KM, KN-M, RO, WRH and SW: methodology, writing, review, and editing. KM, RO, WRH and SW: conceptualization, methodology, formal analysis, supervision, and writing, review, and editing. All authors contributed to the article and approved the submitted version.

## 8 Acknowledgments

We are grateful to Hirokazu Yano for providing valuable information on the Inc18 plasmid. We sincerely thank Yuh Shiwa for supporting ddPCR analysis. We also thank Viktoria Reimann and Alena Kaltenbrunner for providing the information of VIII23 plasmid.

