## Supplementary material for "Discovery of Novel Replication Proteins for Large Plasmids in Cyanobacteria and Their Potential Applications in Genetic Engineering": Supplementary Materials.pdf

#### 1 Supplementary Data

**Supplementary Data 1, Number and list of proteins containing domains typical of CyRepA and CyRepX in cyanobacterial genomes.**

**Supplementary Data 2, Predicted 3D structure data of CyRepX1 (Slr6031)**

**Supplementary Data 3, Predicted 3D structure data of CyRepX2 (Slr6090)**

**Supplementary Data 4, Predicted 3D structure data of A0A132Z0X2**

**Supplementary Data 5, Predicted 3D structure data of Q58352**

**Supplementary Data 6, Complete nucleotide sequence of p6031-1C-GFP.**

**Supplementary Data 7, Complete nucleotide sequence of p6090-1C-GFP.**

**Supplementary Data 8, Complete nucleotide sequence of p6090-ST.**

**Supplementary Data 9, Complete nucleotide sequence of p6090-ST$^{fusion}$.**

### 2 Supplementary Figures and Tables

#### 2.1 Supplementary Figures

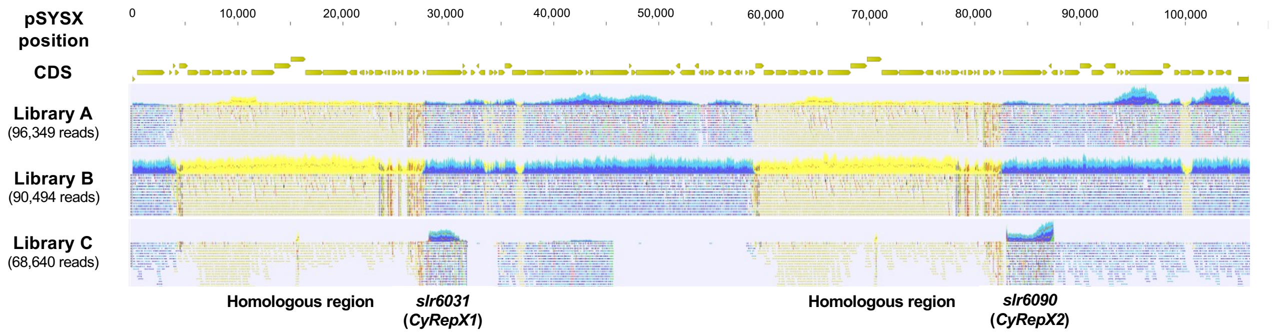


**Supplementary Figure S1. Mapping results of sequence reads obtained from AR-seq to the pSYSX plasmid.**

Reads that were successfully mapped to pSYSX as a pair are shown as blue or light blue, while reads that mapped to multiple regions in the *Synechocystis* 6803 genome are shown in yellow.

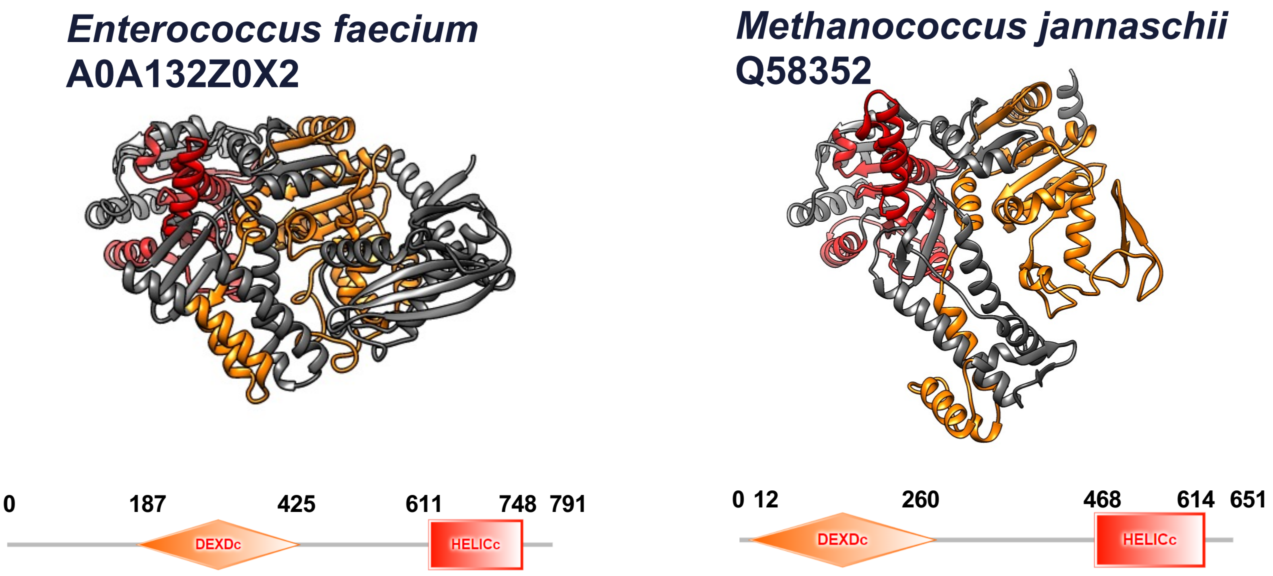


**Supplementary Figure S2. Proteins showing structural similarity to CyRepX.**

3D structures of Rep-related proteins (A0A132Z0X2 and Q58352) predicted using AlphaFold2 were shown along with the domain structure obtained from SMART algorithm. The 3D structures are orange (DEXDc) and red color (HELICc)-coded based on the results of domain predictions by SMART. The numbers indicate the position of the amino acid sequence of each protein.

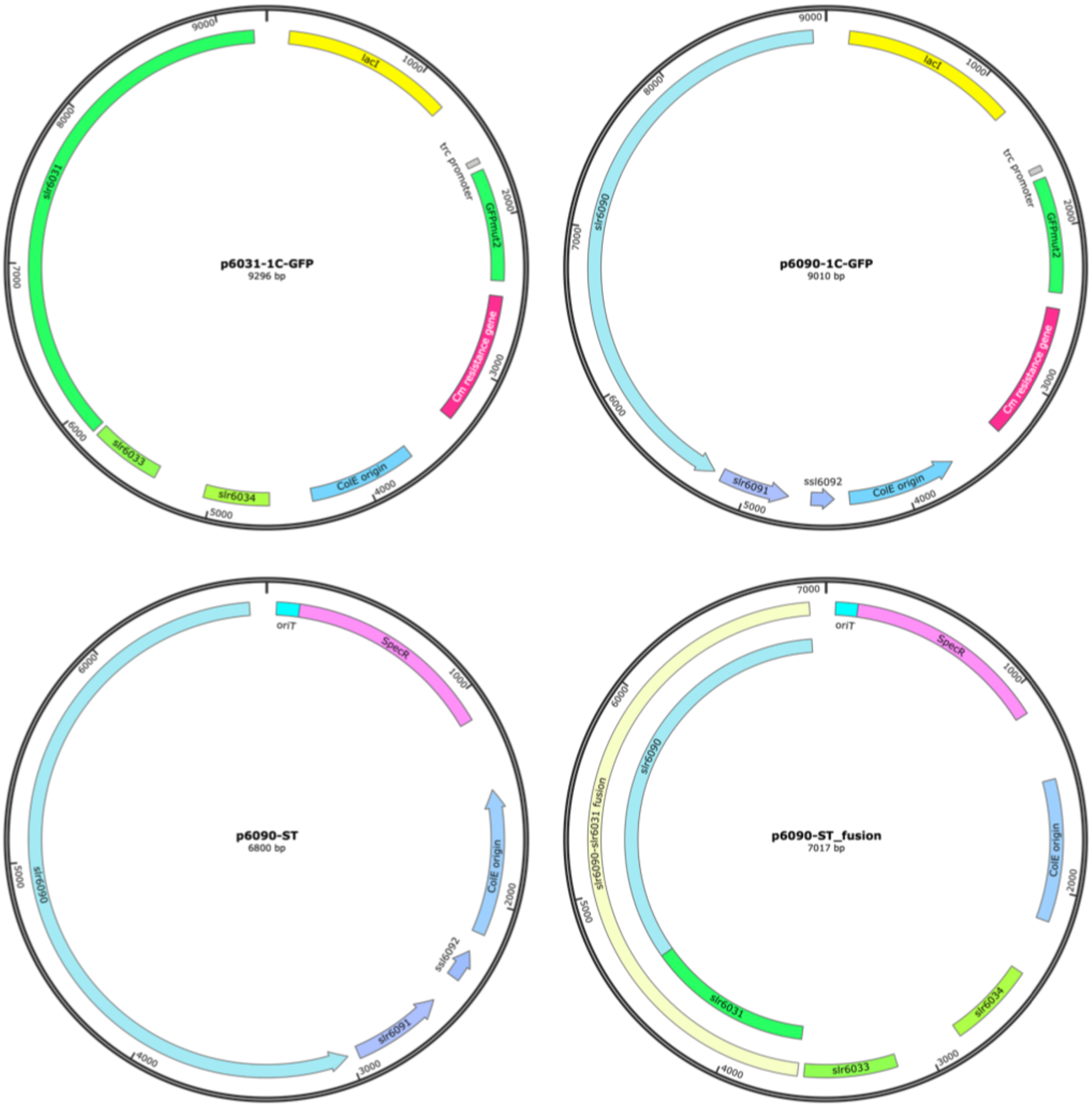


**Supplementary Figure S3. The vector map of p6031-1C-GFP, p6090-1C-GFP and p6031-ST.**

The plasmid vectors p6031-1C-GFP, p6090-1C-GFP and p6090-ST were constructed from plasmids containing the region obtained by screening (*slr6031-slr6034, slr6090-ssl6092* in pSYSX), the *ColE* region and antibiotics resistance marker gene (Cm or Sp). To monitor the expression level in an IPTG-dependent manner, the GFP gene was placed under the *trc* promoter containing the *lacO* operator, together with the repressor *lacI* gene in p6031-1C-GFP and p6090-1C-GFP. The p6090-$ST^{fusion}$ is a derivative obtained by co-introduction of p6031-1C-GFP and p6090-ST in *Synechococcus* 7942, in which part of p6090-ST is replaced by part of p6031-1C-GFP. The image of the vector map was drawn using SnapGene software. The vector sequence can be obtained from the Supplementary Data 6-9.

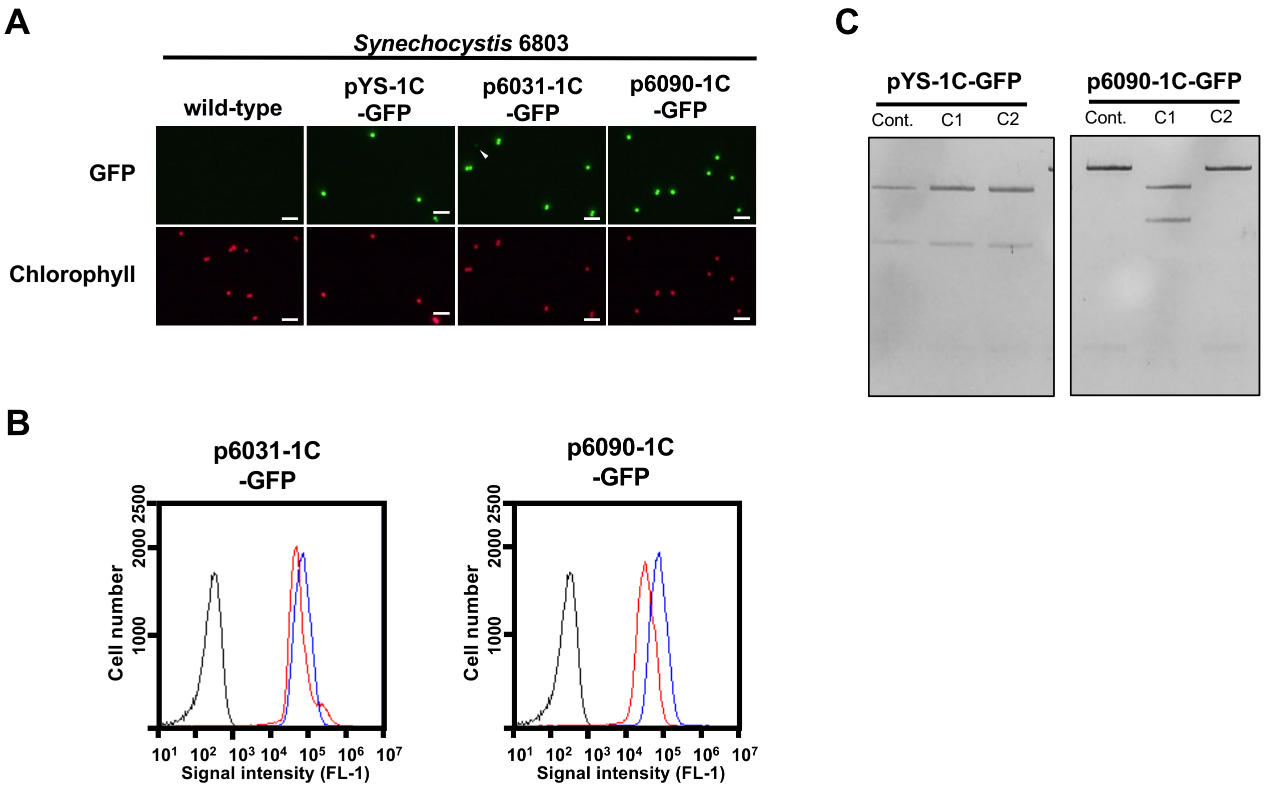


**Figure S4. Analysis of pYS1C-GFP, p6031-1C-GFP, p6090-1C-GFP transformants in *Synechocystis* 6803.**

*Synechocystis* 6803 transformants were cultivated for 2 days. The cells were harvested 2 hours after addition of 1 mM IPTG and compared the GFP and chlorophyll with wild-type strain. (A) Fluorescence microscopy images. White bar:10 µm (B) FACS analysis of GFP fluorescence. Signal intensity of FL1 indicating GFP fluorescence in *Synechocystis* 6803 wildtype (black) pYS1C-GFP (blue) and p6031-1C-GFP or p6090-1C-GFP transformants (red) cultivating with IPTG are shown. (C) Electrophoresis image of plasmids digested by the restriction enzymes, which prepared as shown in Figure 3C. The plasmids before transformation were used as controls. The results of two independent clones are shown as C1 and C2. No plasmid was obtained from the p6031-1C-GFP transformant.

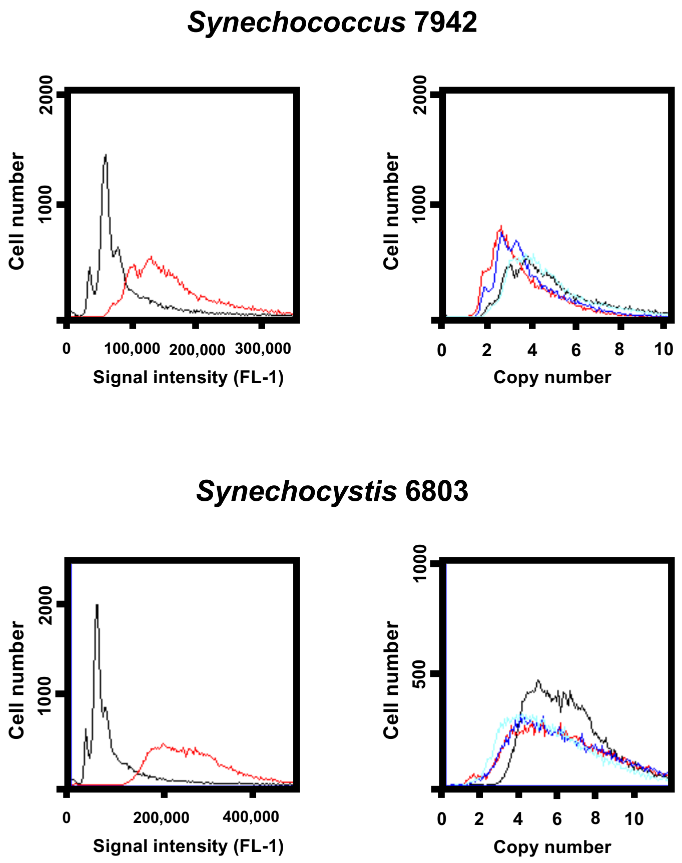


**Figure S5. Estimation of genome copy number.**

After 2 days cultivation under liquid BG-11 medium, *Synechococcus* 7942 and *Synechocystis* 6803 cells were harvested and subjected to the flow cytometry. Left profiles: wild-type (WT) cultures of *Synechococcus* 7942 and *Synechocystis* 6803 (red) and standard sample showing 1-3 copies of *Synechococcus* 7942 cells cultured in phosphate-depleted BG-11 medium for 1 week (Black). Right profiles: Genomic copy number estimated from FL-1 intensity in the left profiles. The copy number of transformants of pYS1C-GFP (red), p6031-1C-GFP (blue), and p6090-1C-GFP (light blue) was shown with WT (black).

#### 2.2 Supplementary Tables

**Supplementary Table S1. Oligonucleotides used in this study.**

| Primer | Sequence (5′ to 3′)[a] |
|---|---|
| *Primers for plasmid construction* | |
| F1 | CTGTCAGACCAAGTTTACTCATATATACTTTAG |
| R2 | CCAGGCATCAAATAAAACGAAAGGCTCAGTCGA |
| F3 | TTATTTGATGCCTGGGGATGCGTATCTAGAAAGAATCGATGTT |
| R4 | AACTTGGTCTGACAGGAAAAAAGAGATTTTTCTGATTTAGT |
| F5 | AAACTTGGTCTGACAGGATCCAAATTTTAAAAAAACCACTCTTTATATCT |
| R6 | TTATTTGATGCCTGGGGATGCGTATCTAGAAATAATCGGTTCT |
| F7 | TTTCGGGGAAATGTGGCCAGCCTCGCAGAGCAG |
| R8 | GTTACCACCGCTGCGGGGCAGGATAGGTGAAGTAGGCC |
| F9 | CGCAGCGGTGGTAACGGCGCAGT |
| R10 | CACATTTCCCCGAAAAGTGCCACCTGA |
| R11 | CCAGGCATCAAATAAAGACGTCAGGTGGCACTTTTCGGGGAAATG |
| *Primers for the estimation of plasmid copy number* | |
| F12 | CCTCCGGTTGGTGGTGGC |
| R13 | GTCCGGCCCTGTTGGGC |
| F14 | CCAAGGTAAGGGTCTTGAGTTTCTTGAC |
| R15 | TCAGCCGTTCAGTCAAGTGAACC |
| F16 | GCCTGATGAATGCTCATCCGGAATTCCGT |
| R17 | GGTTTTCACCGTAACACGCCACATCTTGCG |
| F18 | CCCCTCGTCAAAAATAAGGTTATCAAGTGAGA |
| R19 | AATTGTCCTTTTAACAGCGATCGCGTATTTC |
| *Fluorescent oligo-probes for the estimation of plasmid copy number** | |
| Oligo1 | [HEX]- TGATCCAAGAAGGTAGCCTGGGTTTAGAACG-[BHQ1] |
| Oligo2 | [HEX]- GGTGGATTCGCCAGAGCATGACAAGG-[BHQ1] |
| Oligo3 | [FAM]- CGCTCTGGAGTGAATACCACGACG-[BHQ1] |
| Oligo4 | [FAM]- AACAGGCCAGCCATTACGCTCGTC-[BHQ1] |

*HEX: Hexachlorofluorescein, FAM: Fluorescein, BHQ1: Black Hole Quencher 1

**Supplementary Table S2. Sequencing results of *Synechocystis* 6803 genomic libraries.**

| | Library A | Library B | Library C |
|---|---|---|---|
| chromosome (3,570 kbp) | 412859 (59.390%) | 282098 (30.824%) | 3462 (0.553%) |
| pSYSM (120 kbp) | 25105 (3.611%) | 48575 (5.308%) | 80 (0.013%) |
| **pSYSX (106 kbp)** | **96349 (13.860%)** | **90494 (9.888%)** | **68640 (10.971%)** |
| pSYSA (103 kbp) | 21709 (3.123%) | 56670 (6.192%) | 36 (0.006%) |
| pSYSG (44.3 kbp) | 5256 (0.756%) | 64597 (7.058%) | 1468 (0.235%) |
| pCA2.4 (2.4 kbp) | 8738 (1.257%) | 99816 (10.907%) | 192 (0.031%) |
| pCB2.4 (2.4 kbp) | 3373 (0.485%) | 81557 (8.911%) | 7 (0.001%) |
| pCC5.2 (5.2 kbp) | 121775 (17.517%) | 191390 (20.912%) | 551772 (88.191%) |
| Total | 695164 (100%) | 915197 (100%) | 625657 (100%) |

Sequencing reads in each library were mapped to the *Synechocystis* 6803 genome (chromosome and 7 plasmids, pSYSM, pSYSX, pSYSA, pSYSG, pCA2.4, pCB2.4, and pCC5.2). The number of reads and ratio in each library are shown.
